# Don’t Benchmark Phenomic Prediction Against Genomic Prediction Accuracy

**DOI:** 10.1101/2025.01.09.632209

**Authors:** Fangyi Wang, Mitchell J Feldmann, Daniel E Runcie

## Abstract

Phenomic selection is a new paradigm in plant breeding that uses high-throughput phenotyping technologies and machine learning models to predict traits of new individuals and make selections. This can allow breeders to evaluate more plants in higher throughput more accurately, resulting in faster rates of gain and reduced labor costs. However, phenomic prediction models are frequently benchmarked against genomic prediction models using cross-validation to demonstrate their usefulness to breeders. We argue that this is inappropriate for two reasons: 1) differences in the accuracy statistic measured by cross-validation do not reliably indicate differences in the accuracy parameter of the breeder’s equation, which we show analytically and through re-analysis of data from three representative phenomic prediction studies, and 2) phenomic and genomic selection tools influence other parameters of the breeder’s equation, so comparing accuracy, even if done properly, is insufficient to advocate for one approach over the other. We conclude that phenomic selection may be useful, but comparisons of accuracy between genomic prediction and phenomic prediction models are not.

## 1 INTRODUCTION

Breeders create new varieties by iterating through cycles of population creation, evaluation, and selection to create the next generation. The expected outcome of each cycle is approximated by the breeder’s equation (Lynch et al., 1998):

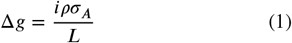

where Δ*g* is the improvement in the average additive genetic value per year; *i* is the standardized selection intensity, the difference in the mean trait value of the selected individuals relative to the whole population, standardized by the standard deviation of trait values; *ρ* is the accuracy of selection, the Pearson correlation coefficient between the estimated and true additive genetic values among the selection candidates; *σ*_*A*_ is the standard deviation of additive genetic values; and *L* is the cycle length, the difference in years between when the population is created and when selections are made. While the Breeder’s equation is only an approximation of the effect of selection in real breeding programs (Piepho and Mohring, 2007; Bijma, 2020), studying how changes in breeding procedures jointly impact these four parameters can give insight into the total impact on genetic gain.

In a traditional breeding program, selection decisions are based on phenotypic measurements of each candidate individual. In many species, collecting this data requires generating lines to grow in field trials and running replicated trials across multiple locations. Therefore, each selection cycle can take several years.

Meuwissen et al. (2001) introduced genomic selection as a tool to increase the rate of gain in breeding programs. Genomic selection means selecting candidate individuals based on estimates of genetic values from a genomic prediction (GP) model using genetic marker data. Genomic selection can increase the rate of gain (Δ*g*) in a breeding program by increasing selection intensity (*i*), increasing selection accuracy (*ρ*), or reducing selection lengths (*L*). If genetic marker data are inexpensive, more candidates can be evaluated using a GP model than can be evaluated directly using field trials. Larger numbers of candidates allow breeders to impose a higher selection intensity while maintaining a constant number of selected individuals to conserve genetic variance (Bernardo et al., 2006; Cobb et al., 2019). GPs of additive genetic values can be more accurate than direct estimates from field trials when candidates are genetically related at causal loci because the sample size per allele can be higher than the sample size per candidate (Zhong et al., 2009). Genomic predictions of additive genetic values can be made as soon as genotype data are collected, even on seeds that have yet to germinate in some species (VanRaden, 2008). This can enable multiple cycles per year (Bhat et al., 2016). However, multiple factors continue to limit the use of genomic selection in plant breeding programs, including the cost of large-scale genotyping, the need to train and continuously update GP models, and the logistics of reorganizing existing breeding schemes.

Rincent et al. (2018) introduced phenomic selection as an alternative tool for breeders that can be less expensive and less logistically challenging than genomic selection. Like genomic selection, phenomic selection also relies on a statistical or machine learning-based model to make selections, called a “phenomic prediction” (PP) model, because instead of genetic markers, the features of the model are measured by high-throughput phenotyping technologies (HTP). For example, Rincent et al. (2018) used multispectral image data from wheat leaves and grains and poplar wood tissue, Winn et al. (2023) used spectral image data from a camera mounted on an aerial vehicle, and Krause et al. (2019) used hyperspectral image data from a camera mounted on an aircraft. Phenomic selection has garnered interest in the breeding community for several reasons: i) HTP data can often be collected at lower cost and higher throughput than genotype data, enabling breeders to evaluate more candidates at once, or evaluate a common set of candidates more times to increase the heritability. ii) Logistically, phenomic selection may require fewer changes to breeding schemes than genomic selection, and many of the same software packages used for implementing GP models also work for PP models. iii) PP models often achieve relative high accuracy when evaluated using cross-validation.

Both genomic and phenomic selection schemes have the potential to accelerate plant breeding. But, since implementing either requires significant investment in equipment and changes to existing breeding schemes, there is considerable interest in determining which approach is best. Multiple recent papers have demonstrated the potential of phenomic selection by showing estimates of accuracies of PP models that are as high or higher than estimates of accuracies of GP models made using the same datasets. However, our objective is to show that these comparisons *are not useful* for evaluating the relative benefits of implementing phenomic and genomic selection schemes in breeding programs.

One of the reasons is that the accuracy statistic reported by these studies is not an appropriate estimator of the selection accuracy parameter of the breeder’s equation (*ρ*). For simplicity, we assume that we have evaluated a set of *n* candidates by some method (*i*.*e*. GP or PP), and that the candidates are unrelated and so independent and identically distributed. While this is not exactly true in any real breeding program, these assumptions are useful for illustrative purposes. The *accuracy* of selecting among the candidates, represented by the parameter *ρ* in the breeder’s equation, is the Pearson correlation coefficient between the estimated (*â*) and true (*a*) breeding values (*i*.*e*. additive genetic values) of the candidates:

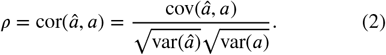

Unfortunately, it is not easy to create a consistent estimator of this parameter because additive genetic values (*a*) cannot be directly observed. Instead, we do observe phenotypes of candidates (*y*). The Pearson correlation coefficient between estimated breeding values and phenotypes (*ρ*_*y*_) is:

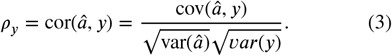

We can easily form a consistent estimator of *ρ*_*y*_ as:

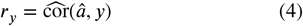

using cross-validation (Legarra et al., 2008; Ould Estaghvirou et al., 2013), where 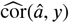) is the sample correlation. Legarra et al. (2008) coined the term *predictive ability* (*ρ*_*y*_) to distinguish it from *accuracy* (*ρ*), but these terms are not used consistently in the plant breeding literature (Winn et al., 2023; Krause et al., 2019). While *r*_*y*_ is a consistent estimator of *ρ*_*y*_, it is not a consistent estimator of *ρ* because *â* and *y* share non-additive genetic and environmental effects, even for individuals not used for model training, as we elaborate on below. Thus, using *r*_*y*_ to predict responses to selection will usually not be accurate.

Another reason is that genomic and phenomic selection schemes impact other parameters of the breeder’s equation differently, so comparisons need to consider the full effects on genetic gain. We will further discuss this in Discussion.

In this study, first, using a more appropriate technique to estimate *accuracy*, we re-analyze published data from Rincent et al. (2018), Winn et al. (2023), and Krause et al. (2019), and show that conclusions about the relative merits of GP and PP change when compared using this method. Then, we demonstrate these observations analytically.

We conclude that phenomic selection may be useful, but comparisons of *accuracies* between GP and PP models are not.

## 2 MATERIALS AND METHODS

We replicated the comparisons between GP and PP *accuracy* reported in three published studies to test how each set of conclusions would change if *accuracies* were estimated using the method in Runcie and Cheng (2019) instead of as *predictive ability*. In each case, we attempted to reproduce the GP and PP models exactly as reported in each original paper.

### 2.1 Case Study 1

Rincent et al. (2018) used two datasets, one from wheat and one from poplar, to compare the *predictive abilities* of GP and PP models.

The wheat data consisted of 223 genotypes measured for heading date and grain yield in two treatments (dry and irrigated). We considered the 4 trait-treatment combinations as 4 traits (heading date in the dry treatment, heading date in the irrigated treatment, grain yield in the dry treatment, and grain weight in the irrigated treatment). Genomic data consisted of 84,529 SNPs, and we used it to train GP models for the 4 traits. Phenomic data consisted of near-infrared spectroscopy scans from both leaf and grain tissues in each treatment. Phenotypic data were reported as best linear unbiased estimators (BLUEs) of genotype means correcting for random block effects. We trained PP model using phenotypes from the 4 traits and their corresponding phenomic data, following the methods in the source paper.

The poplar data consisted of 562 genotypes measured for height, circumference, bud flush, bud set, and rust resistance in two locations: Orleans, France and Savigliano, Italy. All 5 traits were measured at Orleans, but only circumference, bud flush and bud set were measured at Savigliano. Genomic data consisted of 7,808 SNPs. Phenomic data were collected from the wood samples in both locations. Phenotypic data were reported as BLUEs of genotype means correcting for random block effects. We considered each of the 8 trait-location combination as independent traits and used the corresponding phenomic data along with genomic data to train PP and GP models following the methods in the source paper.

For each trait in each species, we fit the following GP model following the methods of Rincent et al. (2018):

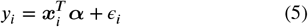

where *y*_*i*_ is the BLUE of the phenotypic value of genotype *i*, ***x***_*i*_ is the SNP markers for genotype 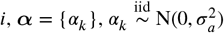 are the random marker effects, and 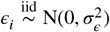 are residuals.

Also following Rincent et al. (2018), we fit the following PP model for each trait in each species:

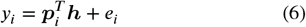

where ***p***_*i*_ is the phenomic features for genotype *i*, ***h*** = {*h*_*k*_}, 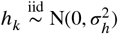 are the random phenomic feature effects, and 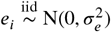 are residuals.

We estimated the *predictive ability* of PP and GP models as the sample Pearson’s Correlation Coefficient between predictions and observed BLUEs using eight-fold cross-validation for the wheat data, and using five-fold cross-validation for the poplar data, following Rincent et al. (2018). We estimated PP *accuracy* using the method by Runcie and Cheng (2019) by fitting a bivariate linear mixed model (see Appendix Part 3) between the observed phenotypic traits and the predicted values from PP models, using the marker-based kinship matrix as the estimate of additive genetic relationsh<u>ips</u> among individuals. We estimated GP accuracy as 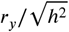 (see Results and Theory Part 3) (Legarra et al., 2008).

### 2.2 Case Study 2

Winn et al. (2023) compared GP and PP *predictive abilities* for grain yield in wheat. The dataset consisted of grain yield data for 3,442 genotypes measured across multiple replicated trials across multiple years and locations. Genotype data consisted of 23,130 SNPs. We removed SNPs with a minor allele frequency of or less than 5%. Phenomic data were derived from images taken by unoccupied aerial vehicles from the same genotypes across different years and different locations. We followed Scenario 1 in Winn et al. (2023) and calculated BLUEs of grain yield and of each spectral band at each genotype, correcting for the random effect of unique combinations of year and location. For GP, we used equation 5 as above, matching the GP model reported by Winn et al. (2023). For PP, we followed the same methods of Winn et al. (2023) and used the following linear model which treats the phenomic features as fixed effects:

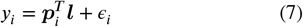

where ***l*** is the vector of fixed effects of the phenomic features. We also used a linear mixed model, equivalent to Equation 6, which treats the phenomic features as mixed effects for PP.

We used five-fold cross-validation to obtain the Pearson’s correlation between the predicted values and the BLUEs. We estimated the *accuracy* in the same way as in Case Study 1.

### 2.3 Case Study 3

Krause et al. (2019) also compared GP and PP *predictive abilities* for grain yield in wheat. The dataset consisted of grain yield data for 3,771 genotypes measured from multiple replicates on the same genotype across different breeding cycles and different management practices. The authors considered each combination of breeding cycle and management practice as a unique treatment, or “site-year” as in Krause et al. (2019). Genomic data were provided as a genomic relationship matrix calculated from 34,900 SNPs. Phenomic data consisted of hyperspectral reflectances collected with a hyperspectral camera mounted in an aircraft. The phenomic data were collected multiple times during the growing season for each treatment. The authors provided the BLUEs of grain yield and of each hyperspectral band for each genotype within each treatment, correcting for random effects of trial, replicate, and block. This was described as the **ALL** method in Krause et al. (2019).

For the grain yield of wheat in Krause et al. (2019), we fit a GP model for genotype *i* within each treatment as follows:

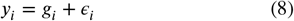

where 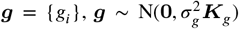 is the random additive genetic effects with genomic relationship matrix ***K***_*g*_ and additive genetic variance parameter 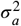. Residuals 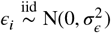 were assumed independent and identically distributed among genotypes. This is the same model as Equation 4 in Krause et al. (2019).

Following Krause et al. (2019), we used the following PP model for genotype *i* within each treatment:

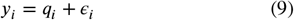

where *y*_*i*_ is the BLUE of phenotypic values of genotype *i* in each treatment, 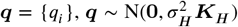 is the random effect of phenomic features on genotype *i* in each treatment with feature covariance matrix 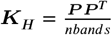, where **P** is the centered and standardized BLUEs of hyperspectral bands of genotype *i* in each treatment, and *nbands* is the total number of hyperspectral bands. This PP model is the same model described in Equation 5 in Krause et al. (2019).

We used five-fold cross-validation to obtain the Pearson’s correlation between the predicted values and the BLUEs within each treatment. We estimated the *accuracy* in the same way as in Case Study 1.

### 2.4 Software

All analyses were conducted in R version 4.2.3 (R Core Team, 2024). Model 7 was fitted using the *lm* function in Base R. Model 5 and 6 were fitted using *mixed*.*solve* function from the *rrBLUP* package (Endelman, 2011). Model 8 and 9 were fitted using the *BGLR* function from *BGLR* package (Pérez and de Los Campos, 2014). *Accuracies* were estimated using the *estimate_gcor* function from the *MegaLMM* package (Runcie et al., 2021) which utilizes *MCMCglmm* package (Hadfield, 2010). BLUEs were calculated using the *lme4* and *emmeans* packages (Bates et al., 2015; Lenth, 2024). Graphs were made with the *ggplot2* and *cowplot* packages (Wickham, 2016; Wilke, 2020).

## 3 RESULTS AND THEORY

### 3.1 Re-analysis of Published Data

We re-analyzed the data from Rincent et al. (2018), Winn et al. (2023), and Krause et al. (2019) to acquire the estimated *predictive abilities* 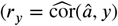 to estimate 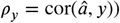 of both GP and PP models using methods mentioned above. Using the method in Runcie and Cheng (2019), we calculated the estimated *accuracies* (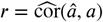 to estimate 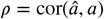) of both GP and PP models, respectively.

For nearly every trait across the three studies, our estimates of the difference between PP and GP *accuracies* were smaller than the reported differences in *predictive abilities*, and all but one points fell below the *y* = *x* line (Figure 1A). Thus, by reporting *predictive abilities*, these studies overestimated the relative benefits of PP. In particular, there were eight traits, including at least one from each of the three studies, where *predictive abilities* were higher for PP, but our estimate of *accuracy* was actually lower for PP, compared to GP. These points are in Quadrant IV of Figure 1A. In all cases, GP *accuracies* were underestimated by *predictive ability* (Figure 1B). In contrast, PP *accuracies* were frequently overestimated by *predictive abilities* (Figure 1B).

**FIGURE 1.**
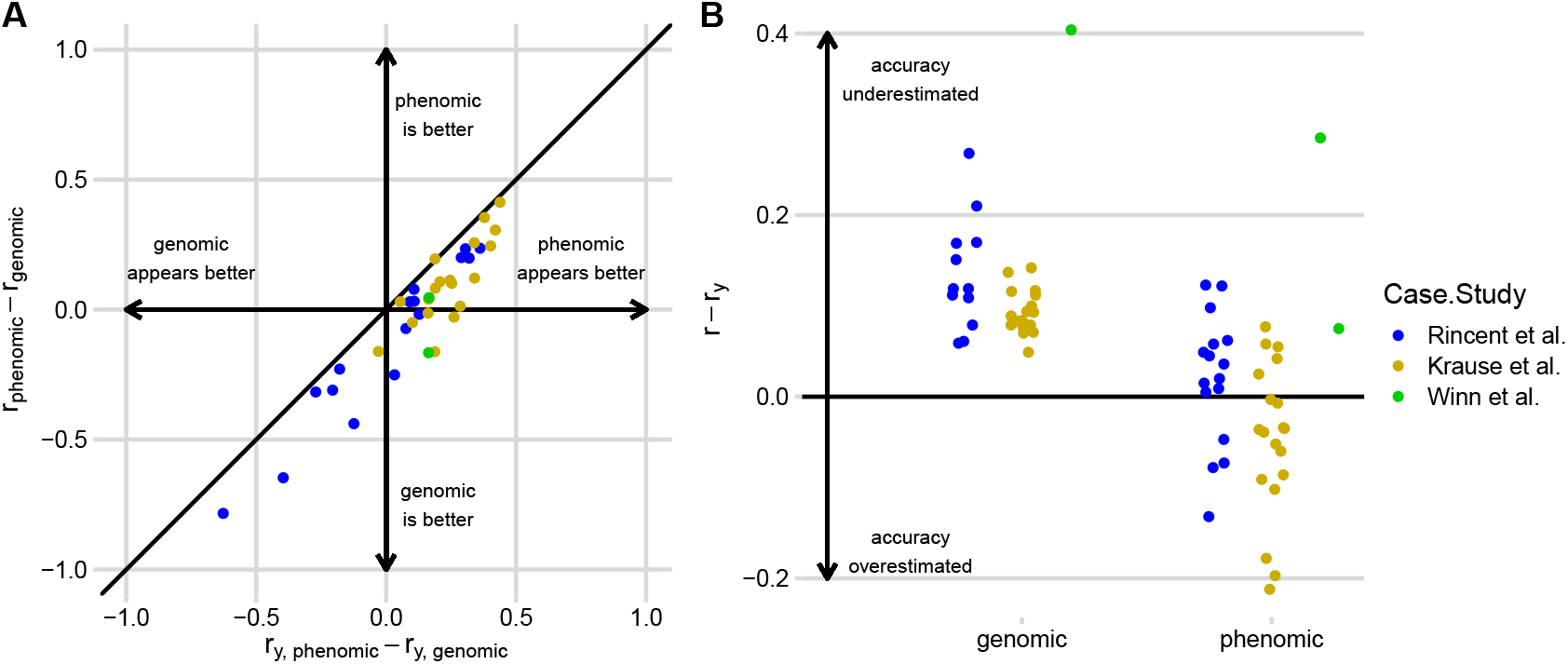
*Predictive ability* overestimates the relative *accuracy* of PP vs GP models. A) Differences between estimated PP and GP *accuracies* (*r*_*phenomic*_ − *r*_*genomic*_ ) (y-axis) are plotted against the differences between estimated PP and GP *predictive abilities* (*r*_*y*,*phenomic*_ − *r*_*y*,*genomic*_ ) (x-axis). The black diagonal shows the 1:1 line. Points to the right of the line indicate that the *accuracy* of PP models tends to be more over-estimated than the *accuracy* of GP models. Points in the lower-right quadrant are cases where PP models appear better than GP models when compared by *Predictive ability* but we estimate that their *accuracy* is actually lower. B) Differences between estimated *accuracies* and *predictive abilities* (*r* − *r*_*y*_) (y-axis) are plotted for GP and PP models, binned by case study. Colors in both panels identify individual traits in each case study.

### 3.2 Sources of bias

We provide simplified analysis and make assumptions for illustrative purposes to demonstrate the sources of bias in the following sections. These assumptions may not be realistic in real-world breeding programs, but we emphasize their importance in conveying our theories.

To understand the source of bias in *r*_*y*_ relative to *ρ*, consider a phenotypic selection scheme on unrelated individuals where *â* = *y*, and a simple path diagram (Lynch et al., 1998; Dekkers, 2007) relating phenotypes (*y*) to additive genetic values (*a*):

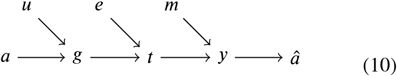

Here, arrows represent causal paths which are assumed to be linear. *a* and *u* are additive and non-additive genetic sources of variation, which together determine the total genetic value (*g*) of a candidate. An organism’s actual trait value *t* is caused by its genetic value plus an effect of its environment (*e*). We make an observation (*y*) which is subject to measurement error (*m*), and use this directly as our selection criterion (*â* = *y*). Based on this diagram, cor(*â, a*) ≠ cor(*â, y*) = 1 because there are exogenous variables that affect *â* = *y* but do not affect *a* (*i*.*e*., *u, e*, and *m*), which we assume are uncorrelated with each other and with *a*. In particular, cov(*â, y*) = var(*y*) ≠ cov(*â, a*) = var(*â*) because *u, e*, and *m* are all common causes of *y* and *â*, but not of *a*.

Next, consider a PP model *f*_*p*_(**p**) = *â*_*p*_ with **p** a vector of phenomic features measured on a candidate. In a phenomic selection scheme, candidates will be ranked by *â*_*p*_, so this is the selection criterion: *â* = *â*_*p*_. To see that *ρ*_*y*_ ≠ *ρ* for *f*_*p*_(**p**), we expand the path diagram to include causes of the phenomic features:

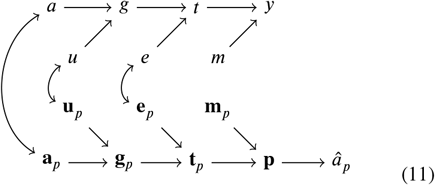

Here, we distinguish the vectors of additive genetic values (**a**_*p*_), total genetic values (**g**_*p*_), and underlying traits (**t**_*p*_) for the phenomic features in **p**, as well as their vectors of non-additive genetic effects (**u**_*p*_), environmental effects (**e**_*p*_), and measurement errors (**m**_*p*_), from the corresponding values and effects underlying *y*. Additionally, various correlations among these effects are possible, the most likely ones of which are drawn with double-headed arrows between the additive genetic effects, non-additive genetic effects, and environmental effects. Thus, even though *f*_*p*_(**p**) = *â*_*p*_ doesn’t involve *y* directly, **p** is caused by exogenous variables **u**_*p*_ and **e**_*p*_ that neither cause *a* nor are correlated with any causes of *a*, but are correlated with causes of *y*. This means that 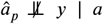, and thus cov(*â*_*p*_, *y*) ≠ cov(*â*_*p*_, *a*).

Finally, consider a GP model *f*_*g*_ (**x**) = *â*_*g*_, with **x** a vector of genotypes for a candidate. We can expand the original path diagram to include the genetic features:

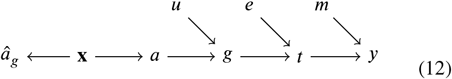

According to this path diagram, *â*_*g*_ ⫫ *y* ∣ *a* because there are no common exogenous variables impacting *â*_*g*_ and *y* that are not also shared with *a*. The only shared exogenous variable is **x** itself. Therefore, cov(*â*_*g*_, *a*) = cov(*â*_*g*_, *y*). However, there are additional sources of variance in *y* that do not affect *a* (*u, e*, and *m*_*y*_), so var(*y*) > var(*a*). Thus, cor(*â*_*g*_, *a*) ≠ cor(*â*_*g*_, *y*).

Note that in this last diagram, we did not include a causal arrow from the genotypes (**x**) to the non-additive genetic effects (*u*). Non-additive genetic effects are caused by genetic features. But additive genetic values and dominance genetic values are uncorrelated (Su et al., 2012; Vitezica et al., 2013, 2016; Ertl et al., 2014; Muñoz et al., 2014; Aliloo et al., 2016, 2017; Xiang et al., 2016). If we use an additive encoding of these features that is orthogonal to dominance coefficients as in Vitezica et al. (2017), and use a GP model that is linear in the genetic features, like rrBLUP (Endelman, 2011), or Bayesian Alphabet models (Gianola, 2013), then we can assume approximately that cov(*f*_*g*_ (**x**), *u*) = 0. In contrast, this independence will not hold for GP models with non-additive encodings of genotype information such as RKHS or Deep Learning models.

### 3.3 Correction factors

In the case of an additive GP model, like rrBLUP, a simple analytical correction is possible to form a consistent estimator of *ρ*: Because cov(*â*_*g*_, *a*) = cov(*â*_*g*_, *y*), *ρ* and *ρ*_*y*_ differ only due to the sources of variance in *y* that are not causes of *a* (*i*.*e*., *u, e*, and *m*_*y*_). Thus, 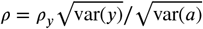, so *r*_*y*_ tends to *underestimate ρ* (Figure 2A). This is the basis for the estimator of *ρ* proposed by Legarra et al. (2008):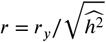, where 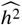 is the estimate of the narrow-sense heritability of *y*.

**FIGURE 2.**
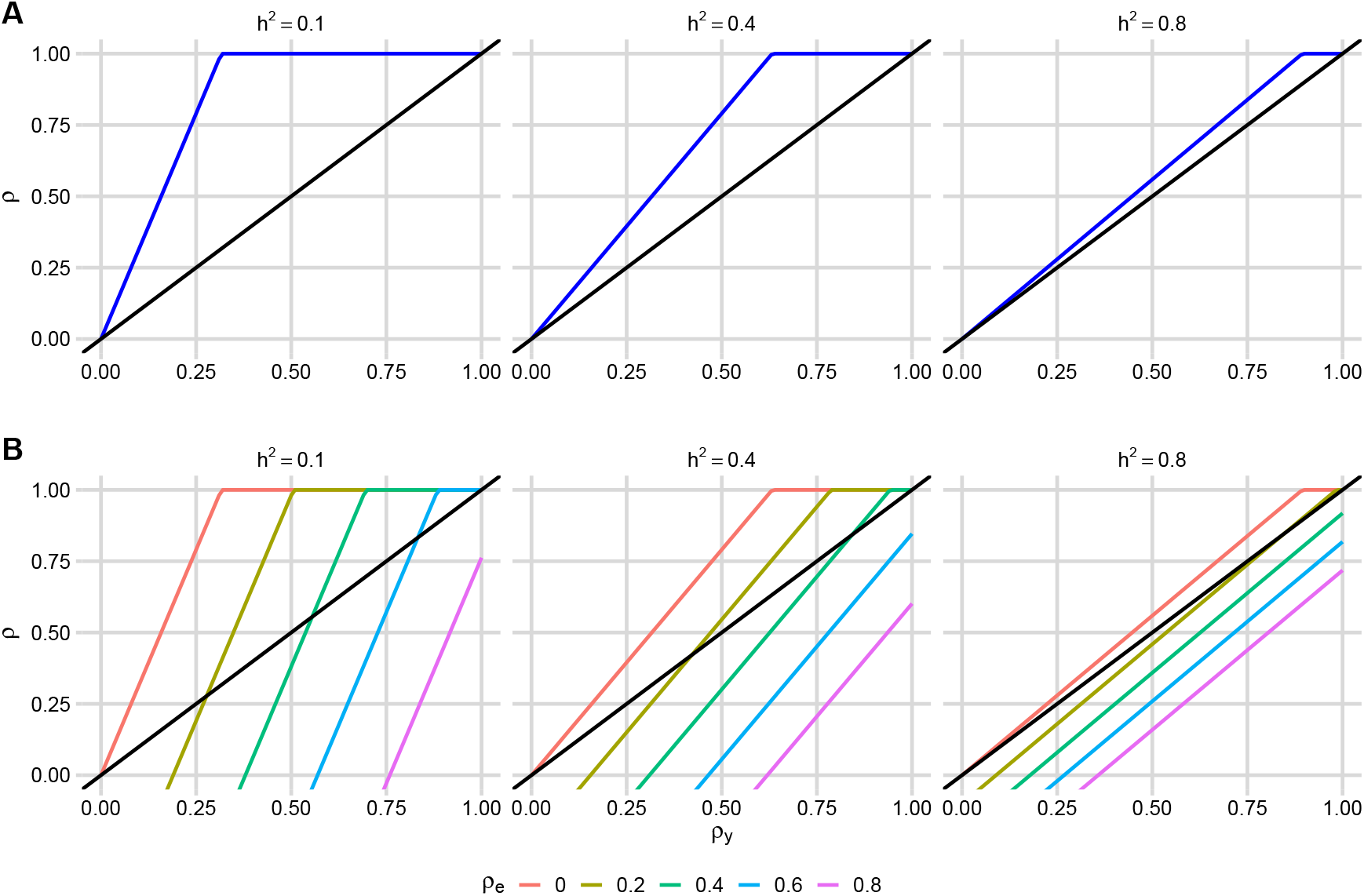
The relationships between *predictive ability* (*ρ*_*y*_) and *accuracy* (*ρ*) for GP (a, blue lines) and PP models (b, colored lines) under 3 levels of heritabilities (0.1, 0.4, 0.8). The black lines indicate *y* = *x*.

This correction factor is not valid for PP because cov(*â*_*p*_, *a*) ≠ cov(*â*_*p*_, *y*) as shown above. Instead, assuming the linear model *y* = *a*+*u*+*e*+*m*_*y*_ and independent candidates, we can show (see Appendix Part 1):

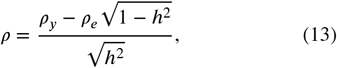

where *ρ*_*e*_ = cor(*â*_*p*_, *u* + *e* + *m*_*y*_). Even if phenomic features (**p**) and the trait (*y*) are not measured on the same plots, so cov(*â, e* +*m*_*y*_) = 0, non-additive genetic variation (*u*) can still cause *ρ*_*e*_ ≠ 0. The impact of *ρ*_*e*_ on the bias of 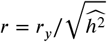 depends on the narrow-sense heritability (*h*^2^), becoming stronger for low-heritability traits. Therefore, cross-validation-based calculations of *r*_*y*_ for a PP model can either underestimate *or overestimate* the actual *accuracy* (Figure 2B).

To demonstrate the impact of *ρ*_*e*_ on *ρ*, we highlight some examples (Figure 2)

1. If the heritability is low, e.g., *h*^2^ = 0.1, and *ρ*_*y*_ = 0.25 for a G<u>P</u> model, its <u>ac</u>tual *accuracy* is much higher: 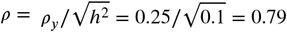. If instead *ρ*_*y*_ = 0.25 for a PP model, the actual *accuracy* could be as high as *ρ* = 0.79 if *ρ*_*e*_ = cor(*â*_*p*_, *u* + *e* + *m*_*y*_) = 0. But this is unlikely unless phenotypes (*y*) and phenomic features **p** are measured on different plots and 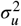 is very small. However, *ρ* could be lower than *ρ*_*y*_ if cor(*â*_*p*_, *u* + *e* + *m*_*y*_) > 0.18. To provide an intuition of *ρ*_*e*_’s relationship with *ρ*, if phenomic features are direct estimates of the trait of interest (*e*.*g*., the trait of interest is plant height and drone images are used to measure plant height instead of a proxy trait), th<u>at is</u>, **a** = *a*, **u** = *u*, **e** = *e* in Diagram 11, then 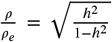 Equation 13 can be simplified to 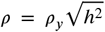 (see Appendix Part 2), and *ρ* would be 0.079 for the PP model.
2. If the heritability is moderate, e.g., *h*^2^ = 0.4, and *ρ*_*y*_ = 0.5 for a G<u>P</u> model, i<u>ts a</u>ctual *accuracy* is much higher: 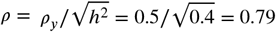. If instead *ρ*_*y*_ = 0.5 for a PP model, the actual *accuracy* could be as high as *ρ* = 0.79, but would equal *ρ* = 0.2 if the correlations of predictions with *a* and *u*+*e*+*m*_*y*_ are equal, and could be as low as *ρ* = 0 if the phenomic data is driven more by environmentally-sensitive traits (so *ρ*_*e*_ is larger). If phenomic features are direct estimates of the trait of interest, *ρ* would be 0.37 for the PP model.
3. If the heritability is high, e.g., *h*^2^ = 0.8, differences between *ρ*_*y*_ and *ρ* are smaller for both GP and PP models. But even if *ρ*_*y*_ is very high for a PP model (*e*.*g ρ*_*y*_ = 0.8), the actual *accuracy* can be *ρ* < 0.6, and would be only *ρ* = 0.72 if phenomic features were direct estimates of the trait of interest.

While the difference between *ρ*_*y*_ and *ρ* have been known for many years (Legarra et al., 2008; Ould Estaghvirou et al., 2013), many studies continue to report *r*_*y*_ values when comparing GP models. Despite *r*_*y*_ tending to underestimate the actual *accuracy*, comparing *r*_*y*_ values between two (additive) GP models (say model (1) is RRBLUP and model (2) is BayesC*π*) is valid *if the comparison is based on the ratio* 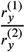 (Legarra et al., 2008):

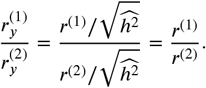

Therefore, the conclusion of the superiority of one model over the other would be the same whether *r*_*y*_ or *r* were used. The same is not true if models are compared based on the *difference* of *r*_*y*_ estimates:

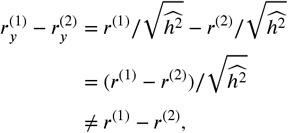

although the *sign* of the comparison remains the same. However, as shown in Figure 2, if a GP and PP model are compared either as a ratio 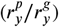 or a difference 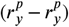, both the sign and the magnitude of the result can differ from the true values (*ρ*^*p*^/*ρ*^*g*^ or *ρ*^*p*^ − *ρ*^*g*^, respectively). This means that conclusions about the superiority of PP over GP based on comparisons of *predictive ability* (*ρ*_*y*_) are unreliable.

### 3.4 Alternative estimates of accuracy

Legarra and Reverter (2018) proposed an alternative estimator of *accuracy* called the “LR method”. Unfortunately this method is not appropriate for PP models because the method assumes that cor(*â, u* + *e*) = 0 which is generally violated for PP models, as discussed above.

Runcie and Cheng (2019) demonstrated three alternative measures of *accuracy* more appropriate for PP models, or models that combined phenomic and genomic features. Of these, the “parametric” approach is the most generally applicable, although it requires genetic or pedigree relationships among testing individuals to compute. Specifically, a consistent estimator of *ρ* can be calculated as (Lopez-Cruz et al., 2020):

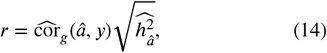

where 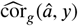 is an estimate of the genetic correlation between predictions (*â*) and phenotypes (*y*) from a bivariate linear mixed model (see Appendix Part 3) fit to testing candidates not used for model training, and 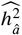 is the estimated heritability of *â*. Large sets of testing candidates are needed to estimate this parameter accurately. We used this method above to re-estimate the *accuracies* of PP models using data from three published studies because it is easier to estimate than equation 13.

## 4 DISCUSSION

Our main goal with this article is to clearly demonstrate that comparing *predictive ability* estimates (*r*_*y*_, the Pearson correlation coefficient between predicted genetic values and observed traits) between PP and GP models is not an appropriate way to assess the relative utility of phenomic selection and genomic selection for breeding programs.

As we show analytically and using previously published empirical results, the *predictive ability* is a biased and inconsistent estimate of the *accuracy* of genetic values (*ρ*), a key parameter of the Breeder’s equation that determines the genetic gain after selection. While the *predictive ability* of an (additive) GP model is always less than its *accuracy*, the *predictive ability* of a PP model can be greater than its *accuracy*. For example, under reasonable, though simplified, conditions, a *predictive ability* of *ρ*_*y*_ = 0.5 for a GP model can indicate an *accuracy* of *ρ* = 0.79 (with *h*^2^ = 0.4). But under the same conditions, a *predictive ability* of *ρ*_*y*_ = 0.5 for a PP model might indicate an *accuracy* of *ρ* = 0.2 (if the correlations of predictions with *a* and *u* + *e* + *m*_*y*_ are equal), or even lower (if predictions are dominated by non-genetic sources of variation). Therefore, comparisons of *predictive abilities* between GP and PP models will tend to give the false impression that PP models are more accurate, even when they are not. Note that in real breeding programs, Equation 13, and the Breeder’s equation itself, may not be strictly appropriate due to correlations among selection candidates and correlated errors due to fixed effects and spatial effect terms in GP or PP models (Piepho and Mohring, 2007; Bijma, 2020). Equation 14 by Runcie and Cheng (2019) generally offers a better solution to estimate PP *accuracy*, which we have used in our results, but doesn’t account for effects of correlations among candidates in calculations of selection intensity. Nevertheless, these simplified scenarios help intuitively demonstrate the reason that *preditive ability* fails as a fair comparison statistic between PP and GG models. We also presented examples of comparisons between PP and GP models from the recent literature, using the same datasets to directly estimate *accuracy* using an alternative method, and show that the conclusions about the superiority of one method over the other frequently change when the more appropriate statistic is used to compare models.

However, even if appropriate methods are used to estimate *accuracyies* (*ρ*) for both GP and PP models, comparing these *accuracy estimates* directly is still not sufficient to compare genomic and phenomic selection breeding schemes. Breeding schemes should be evaluated based on their expected genetic gains over a specific time interval, and using a specific budget. Comparisons between estimates of *ρ* alone are insufficient for this because genomic and phenomic selection schemes are likely to impact the other three parameters of the Breeder’s equation differently, which can overcome differences in accuracy. The other three parameters of the breeder’s equation are *L*, the cycle length, *σ*_*A*_, the additive genetic standard deviation, and *i*, the standardized selection intensity.

The most consequential parameter in the Breeder’s equation is generally *L*, the cycle length. In many breeding programs, each cycle takes multiple years. For example, in Bhat et al. (2016); Tessema et al. (2020) (wheat), the cycle is ≈ 6−7 years. In the University of California strawberry breeding program, we estimated the average cycle length to be 10 years (Pincot et al., 2021). In its most aggressive form, genomic selection can be used to shorten breeding cycles to ≪ 1 year, using the two-part GS scheme proposed by Gaynor et al. (2017), in which candidates are selected based on seed or seedling genotypes without any phenotypic evaluation. This strategy is generally not possible for phenomic selection because most PP models require plants to be grown to mature stages to collect phenomic data. Most apparently successful PP models for yield and quality-related traits use features collected in field trials of candidates close in time to when yield measures are taken (Winn et al., 2023; Krause et al., 2019). Applied this way, phenomic selection would not shorten cycle lengths significantly. Thus, a genomic selection scheme like that proposed by Gaynor et al. (2017) would be preferred over a phenomic selection scheme even if the GP *accuracy* was lower than the PP *accuracy*.

The aggressive two-part GS scheme proposed by Gaynor et al. (2017) is not currently widely used. More commonly, GS only slightly shortens breeding cycles, for example, using GP to select at the F5 or PYT stages of a wheat breeding scheme instead of the AYT stage (Tessema et al., 2020). In this case, differences in *L* may not be large between a genomic selection and phenomic selection scheme. However, one of the principle arguments for phenomic selection is that is that phenomic data can be collected more cheaply and in higher throughput than genomic data. If so, and if each field plot is sufficiently inexpensive, PPs could be made on much larger numbers of candidates, enabling breeders to increase selection intensity. The relationship between population size and selection intensity can be approximated as:

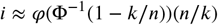

where *n* is the number of candidates, *k* is the target number of selections, and *φ*(.) is the density function and Φ^−1^(.) is the inverse of the cumulative density function of the standard normal distribution. Figure 3 shows that *i* is a function of both *n* and *k*/*n*, the proportion selected. Larger *n* increases *i* but at a relatively slow rate, as discussed by Cobb et al. (2019). Increasing the number of candidates from GP to PP two-fold (*i*.*e*., 100%), from *n*_*g*_ = 100 for a genomic selection scheme to *n*_*p*_ = 200 for a phenomic selection scheme, only increases selection intensity ≈ 25%. Increasing the size of the phenomic selection scheme 10-fold (*i*.*e*., 900%) to *n* = 1, 000) increases intensity ≈ 70% (Figure 3). Both numbers decreases if the genomic selection scheme is larger (*e*.*g. n*_*g*_ = 1, 000, compared to phenomic selection schemes of *n*_*p*_ = 2, 000 or *n*_*p*_ = 10, 000, respectively). Therefore, exponentially larger numbers of candidates must be evaluated for the increase in *i* enabled by phenomic selection to compensate for decreased *accuracy*. Such massive population sizes are often infeasible. Population sizes can be limited by field space and field costs, so it may not be common for the number of selection candidates to be significantly limited when applying genomic selection. Therefore, phenomic selection schemes are unlikely to have a major advantage over genomic selection solely based on increased intensity.

**FIGURE 3.**
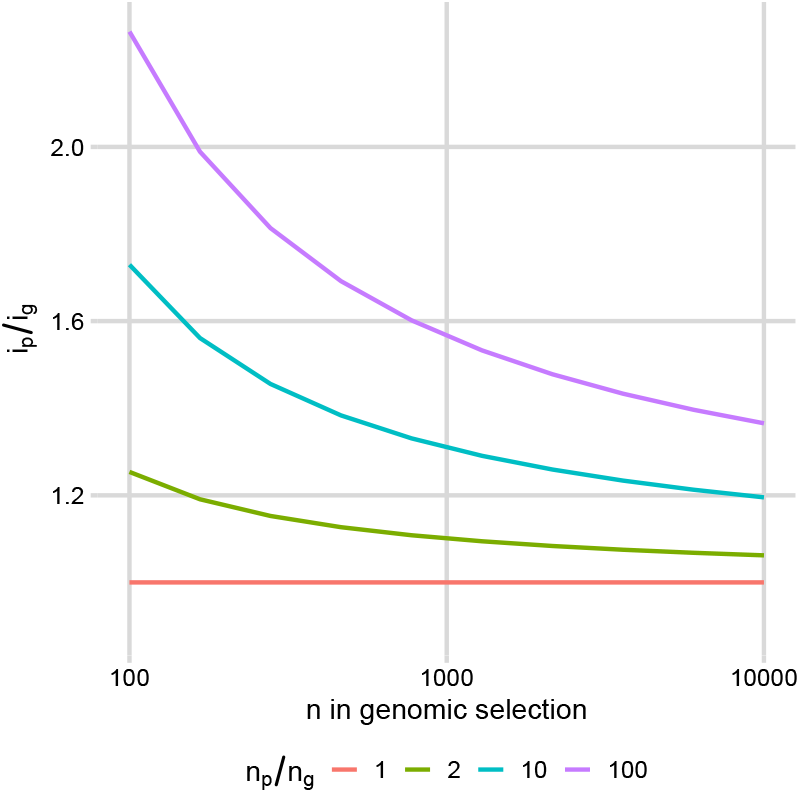
Relationship between selection intensity and population size.

The effects of genomic selection and phenomic selection on the maintenance of genetic variance 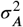 are less clear than for the other parameters of the Breeder’s equation. Jannink (2010) and Goddard et al. (2011) argued that genomic selection should maintain genetic variance longer than pedigree-based selction because genetic markers can be used to select individuals with good alleles at QTL but which are otherwise unrelated. However, others have argued that genomic selection can rapidly reduce genetic variance, and multiple strategies have been suggested to mitigate against rapid genetic variance loss during genomic selections. Phenomic selection is really a form of phenotypic selection, thus the consequences of phenomic selction on genetic variance are likely similar. In either strategy, careful management of genetic variance will be necessary to maintain long-term gains.

Lastly, comparisons of PP and GP predictive ability generally use training datasets that are the same size in an attempt to make the comparison fair. However, this choice itself likely biases the comparison between these approaches. Even if *r* could be estimated accurately, the actual *accuracy* using either strategy will depend on the size and composition of the training data for each model. Since phenomic data are generally cheaper to acquire than genomic data, an optimized phenomic selection scheme would likely permit larger training data sets than an optimized genomic selection scheme. Therefore, PP and GP models should be compared only when each model is trained on an appropriately sized training dataset.

As a final note, our discussions throughout assume that the Breeder’s equation is sufficient to predict genetic gain, and thus that accuracy (*ρ*) is proportional to the rate of genetic gain. But, as discussed by Bijma (2020), this is only exact when the selection index (here: *â*) and breeding values are linearly related (*e*.*g*., bivariate normal). In cases of extreme departures from this assumption, it is mathematically possible for a model with a lower accuracy (*ρ*) to lead to faster gain than a model with higher accuracy, controlling for *i, σ*_*a*_, and *L*. In these cases, the Price Equation, or Robertson–Price identity, will better predict the rate of genetic gain. Nevertheless, these conditions are likely fairly unusual, and we believe that if used carefully, accuracy can still be a useful statistic for studying changes to breeding schemes.

Our goal is not to suggest that phenomic selection cannot be useful for breeding. When PP is used to increase the precision or throughput of phenomic evaluation of candidates, this can certainly improve breeding outcomes. In particular, if PP is used to improve the phenotype data available to train GP models, this can greatly improve breeding outcomes. In this application, it is actually the *predictive ability* of a PP model that matters - the ability of that model to produce highaccuracy replacements for the trait data needed to train an accurate GP model. Therefore, we suggest that PP studies should report *predictive ability*, while GP studies should report *accuracy*. But these two numbers should not be directly compared - they measure different things.

## 5 CONCLUSIONS

To summarize, we argue that comparisons between *predictive abilities* of GP and PP models are largely uninformative for breeders. These comparisons use a statistic that is not directly relevant to breeding goals, and neglect the impact of each strategy on other equally important consequences of the implementation of genomic and phenomic selection schemes.

## Supporting information

Supplemental Table 1: mean and SE of prediction accuracies and predictive abilities

## 6 DATA AVAILABILITY

Scripts used in this study are available at https://github.com/Faye-Wong-stat/breeder_equation

## 7 ACKNOWLEDGEMENTS

The authors would like to acknowledge helpful comments from Hans-Peter Piepho and feedback from Renaud Rincent and Zachary Winn. UC Davis is located at Davis, California, which is located on land that was the home of the Patwin people for thousands of years.

## 8 CONFLICT OF INTEREST

The authors declare that the research was conducted without any commercial or financial relationships that could be construed as a potential conflict of interest.

## 9 AUTHOR CONTRIBUTIONS

FW, DER, and MJF contributed to the conception of this project. FW performed simulations and analyses. FW, DER, MJF wrote and edited the article.

## 10 APPENDIX

### 10 Derivation of correction factor for *ρ*_*y*_

Let *f* (**w**) = *ŷ* be a prediction model that takes the feature vector **w**. Assuming the following decomposition of *y* into additive (*a*), nonadditive genetic (*u*) and residual (*e*) variation (where we combine *e* and *m*_*y*_ into the same term for simplicity):

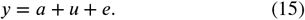

We find:

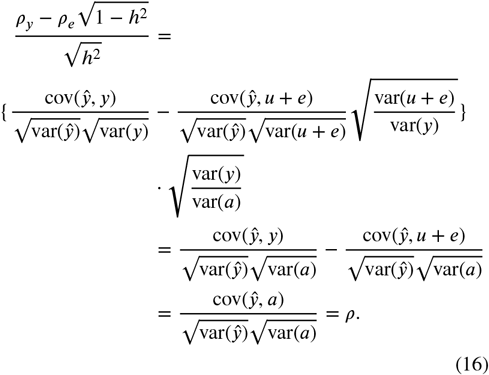

If we can assume that *ρ*_*e*_ = cor(*ŷ, u*+*e*) = <u>0,</u> then the quantity on the left simplifies to the familiar 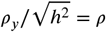. This is often a reasonable assumption for additive genomic prediction models, but is not a reasonable assumption for any phenomic prediction model because cor(*ŷ, u*) ≠ 0. Furthermore, whenever the features **w** are measured on the same individuals as *y*, cor(*ŷ, e*) ≠ 0.

Runcie and Cheng (2019) provided a similar “semi-parametric accuracy estimate” for a prediction model *g*(**x, w**) that takes in both genetic (**x**) and phenomic (**w**) features. This estimate also accounts for non-independence among the selection candidates through a genomic relationship matrix **K**.

### 10.2 Derivation of *ρ*_*e*_ when PP measures *y* directly

If phenomic features are measurements of the targeted trait, for example, plant height, and we assume there is little measurement error on the phenotype (*i*.*e. m*_*y*_ becomes 0) but some measurement error on the phenomic prediction, then Diagram 11 becomes:

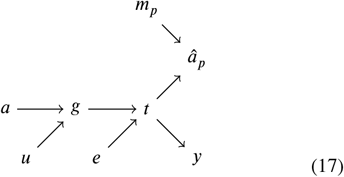

Here, we use *â*_*p*_ instead of **P** for simplification. In the simplest case, *y* = *a* + *u* + *e* and *â*_*p*_ = *a* + *u* + *e* + *m*_*p*_. *ρ* and *ρ*_*e*_ become:

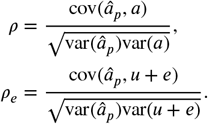

and 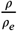 becomes:

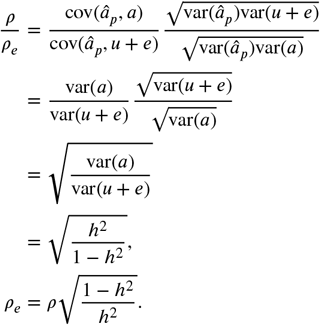

We plug this into Equation 13:

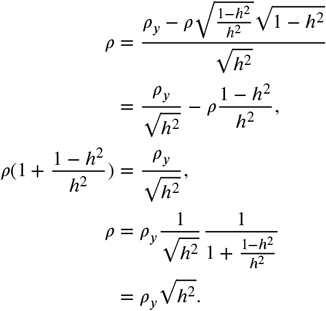

### 10.3 Derivation of estimating accuracy with bivariate linear mixed model

In PP, we can consider **Y** = [**y**, *â*_*p*_] an *n* × 2 matrix. Although we calculated *â*_*p*_ using phenomic data, we assume that it can still be written as an additive linear model with genetic (**a**_*p*_) and environmental (**e**_*p*_) components, and we can write a bivariate linear model as following:

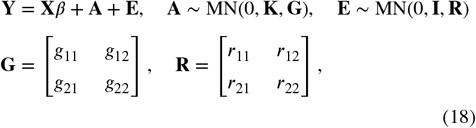

where **X** is a *n* × 2 matrix of 1’s, and *β* a 2 × 1 vector of intercepts, **A** = [**a, a**_*p*_] and **E** = [**e, e**_*p*_], *MN*(⋅, ⋅, ⋅) is the matrix normal distribution. By fitting Model 18, we can estimate 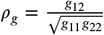 and 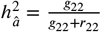, and plug these into equation 14 to estimate PP accuracy. In the analyses in the main text we fit the model with *estimate_gcor()* function from https://github.com/deruncie/MegaLMM_analyses/blob/master/Wheat/Estimate_gcor_prediction.R which uses a Gibbs sampler implemented in the *MCMCglmm* function of the *MCMCglmm* package (Hadfield, 2010) with the following parameters:

~~~
prior = list(R = list(V = diag(c(.5,.01),2),nu = 3),
       G = list(G1 = list(V = diag(c(.5,.5),2),
                nu = 3,
                alpha.mu = rep(0,2),
                alpha.V = diag(1,2))))
nitt = 30000
thin = 50
burn = 5000
~~~

to approximate a flat prior on *ρ*_*g*_ . For **y**, we pre-processed all phenotype data at once and then split the processed data into testing and training partitions for cross-validation.

## Notes

### Competing Interest Statement

The authors have declared no competing interest.

### Summary of Updates

Theory has been moved to the results section. We changed the order of Figure 1 and 2, and also revised Figure 1. We moved the table to the supplementary material section. We re-analyzed data from case study 3. We added 2 parts to the appendix to better explain our analysis.

## REFERENCES

Aliloo, H., Pryce, J. E., González-Recio, O., Cocks, B. G., Goddard, M. E. and Hayes, B. J. (2017) Including nonadditive genetic effects in mating programs to maximize dairy farm profitability. Journal of Dairy Science, 100, 1203–1222. URL: https://www.journalofdairyscience.org/article/S0022-0302(16)30834-7/fulltext. Publisher: Elsevier.

Aliloo, H., Pryce, J. E., González-Recio, O., Cocks, B. G. and Hayes, B. J. (2016) Accounting for dominance to improve genomic evaluations of dairy cows for fertility and milk production traits. Genetics Selection Evolution, 48, 8. URL: 10.1186/s12711-016-0186-0.

Bates, D., Mächler, M., Bolker, B. and Walker, S. (2015) Fitting linear mixed-effects models using lme4. Journal of Statistical Software, 67, 1–48.

Bernardo, R., Moreau, L. and Charcosset, A. (2006) Number and Fitness of Selected Individuals in Marker-Assisted and Phenotypic Recurrent Selection. Crop Science, 46, 1972–1980. URL: https://onlinelibrary.wiley.com/doi/abs/10.2135/cropsci2006.01-0057. _eprint: https://onlinelibrary.wiley.com/doi/pdf/10.2135/cropsci2006.01-0057.

Bhat, J. A., Ali, S., Salgotra, R. K., Mir, Z. A., Dutta, S., Jadon, V., Tyagi, A., Mushtaq, M., Jain, N., Singh, P. K., Singh, G. P. and Prabhu, K. V. (2016) Genomic Selection in the Era of Next Generation Sequencing for Complex Traits in Plant Breeding. Frontiers in Genetics, 7, 221. URL: https://www.ncbi.nlm.nih.gov/pmc/articles/PMC5186759/.

Bijma, P. (2020) The Price equation as a bridge between animal breeding and evolutionary biology. Philosophical Transactions of the Royal Society B: Biological Sciences, 375, 20190360. URL: https://royalsocietypublishing.org/doi/10.1098/rstb.2019.0360. Publisher: Royal Society.

Cobb, J. N., Juma, R. U., Biswas, P. S., Arbelaez, J. D., Rutkoski, J., Atlin, G., Hagen, T., Quinn, M. and Ng, E. H. (2019) Enhancing the rate of genetic gain in public-sector plant breeding programs: lessons from the breeder’s equation. TAG. Theoretical and Applied Genetics. Theoretische Und Angewandte Genetik, 132, 627–645. URL: https://www.ncbi.nlm.nih.gov/pmc/articles/PMC6439161/.

Dekkers, J. (2007) Prediction of response to marker-assisted and genomic selection using selection index theory. Journal of animal breeding and genetics, 124, 331–341.

Endelman, J. B. (2011) Ridge regression and other kernels for genomic selection with r package rrblup. The plant genome, 4.

Ertl, J., Legarra, A., Vitezica, Z. G., Varona, L., Edel, C., Emmerling, R. and Götz, K.-U. (2014) Genomic analysis of dominance effects on milk production and conformation traits in Fleckvieh cattle. Genetics Selection Evolution, 46, 40. URL: 10.1186/1297-9686-46-40.

Gaynor, R. C., Gorjanc, G., Bentley, A. R., Ober, E. S., Howell, P., Jackson, R., Mackay, I. J. and Hickey, J. M. (2017) A two-part strategy for using genomic selection to develop inbred lines. Crop Science, 57, 2372–2386.

Gianola, D. (2013) Priors in Whole-Genome Regression: The Bayesian Alphabet Returns. Genetics, 194, 573–596. URL: https://www.ncbi.nlm.nih.gov/pmc/articles/PMC3697965/.

Goddard, M. E., Hayes, B. J. and Meuwissen, T. H. E. (2011) Using the genomic relationship matrix to predict the accuracy of genomic selection. Journal of Animal Breeding and Genetics, 128, 409–421.

Hadfield, J. D. (2010) Mcmc methods for multi-response generalized linear mixed models: The MCMCglmm R package. Journal of Statistical Software, 33, 1–22. URL: https://www.jstatsoft.org/v33/i02/.

Jannink, J.-L. (2010) Dynamics of long-term genomic selection. Genetics Selection Evolution, 42, 35. URL: 10.1186/1297-9686-42-35.

Krause, M. R., González-Pérez, L., Crossa, J., Pérez-Rodríguez, P., Montesinos-López, O., Singh, R. P., Dreisigacker, S., Poland, J., Rutkoski, J., Sorrells, M., Gore, M. A. and Mondal, S. (2019) Hyperspectral Reflectance-Derived Relationship Matrices for Genomic Prediction of Grain Yield in Wheat. G3 Genes|Genomes|Genetics, 9, 1231–1247. URL: https://doi.org/10.1534/g3.118.200856.

Legarra, A. and Reverter, A. (2018) Semi-parametric estimates of population accuracy and bias of predictions of breeding values and future phenotypes using the lr method. Genetics Selection Evolution, 50, 1–18.

Legarra, A., Robert-Granié, C., Manfredi, E. and Elsen, J.-M. (2008) Performance of genomic selection in mice. Genetics, 180, 611–618.

Lenth, R. V. (2024) emmeans: Estimated Marginal Means, aka Least-Squares Means. URL: https://CRAN.R-project.org/package=emmeans. R package version 1.10.5.

Lopez-Cruz, M., Olson, E., Rovere, G., Crossa, J., Dreisigacker, S., Mondal, S., Singh, R. and Campos, G. d. l. (2020) Regularized selection indices for breeding value prediction using hyper-spectral image data. Scientific Reports, 10, 8195.

Lynch, M., Walsh, B. et al. (1998) Genetics and analysis of quantitative traits, vol. 1. Sinauer Sunderland, MA.

Meuwissen, T. H. E., Hayes, B. J. and Goddard, M. E. (2001) Prediction of Total Genetic Value Using Genome-Wide Dense Marker Maps. Genetics, 157, 1819–1829. URL: 10.1093/genetics/157.4.1819.

Muñoz, P. R., Resende, Jr, M. F. R., Gezan, S. A., Resende, M. D. V., de los Campos, G., Kirst, M., Huber, D. and Peter, G. F. (2014) Unraveling Additive from Nonadditive Effects Using Genomic Relationship Matrices. Genetics, 198, 1759–1768. URL: 10.1534/genetics.114.171322.

Ould Estaghvirou, S. B., Ogutu, J. O., Schulz-Streeck, T., Knaak, C., Ouzunova, M., Gordillo, A. and Piepho, H.-P. (2013) Evaluation of approaches for estimating the accuracy of genomic prediction in plant breeding. BMC Genomics, 14, 860. URL: https://www.ncbi.nlm.nih.gov/pmc/articles/PMC3879103/.

Pérez, P. and de Los Campos, G. (2014) Genome-wide regression and prediction with the bglr statistical package. Genetics, 198, 483–495.

Piepho, H.-P. and Mohring, J. (2007) Computing heritability and selection response from unbalanced plant breeding trials. Genetics, 177, 1881–1888. URL: 10.1534/genetics.107.074229.

Pincot, D. D. A., Ledda, M., Feldmann, M. J., Hardigan, M. A., Poorten, T. J., Runcie, D. E., Heffelfinger, C., Dellaporta, S. L., Cole, G. S. and Knapp, S. J. (2021) Social network analysis of the genealogy of strawberry: retracing the wild roots of heirloom and modern cultivars. G3 Genes|Genomes|Genetics, 11, jkab015. URL: 10.1093/g3journal/jkab015.

R Core Team (2024) R: A Language and Environment for Statistical Computing. R Foundation for Statistical Computing, Vienna, Austria. URL: https://www.R-project.org/.

Rincent, R., Charpentier, J.-P., Faivre-Rampant, P., Paux, E., Le Gouis, J., Bastien, C. and Segura, V. (2018) Phenomic selection is a low-cost and high-throughput method based on indirect predictions: proof of concept on wheat and poplar. G3: Genes, Genomes, Genetics, 8, 3961–3972.

Runcie, D. and Cheng, H. (2019) Pitfalls and remedies for cross validation with multi-trait genomic prediction methods. G3: Genes, Genomes, Genetics, 9, 3727–3741.

Runcie, D. E., Qu, J., Cheng, H. and Crawford, L. (2021) Megalmm: mega-scale linear mixed models for genomic predictions with thousands of traits. Genome biology, 22, 1–25.

Su, G., Christensen, O. F., Ostersen, T., Henryon, M. and Lund, M. S. (2012) Estimating Additive and Non-Additive Genetic Variances and Predicting Genetic Merits Using Genome-Wide Dense Single Nucleotide Polymorphism Markers. PLOS ONE, 7, e45293. URL: https://journals.plos.org/plosone/article?id=10.1371/journal.pone.0045293. Publisher: Public Library of Science.

Tessema, B. B., Liu, H., Sørensen, A. C., Andersen, J. R. and Jensen, J. (2020) Strategies Using Genomic Selection to Increase Genetic Gain in Breeding Programs for Wheat. Frontiers in Genetics, 11, 578123. URL: https://www.ncbi.nlm.nih.gov/pmc/articles/PMC7748061/.

VanRaden, P. M. (2008) Efficient Methods to Compute Genomic Predictions. Journal of Dairy Science, 91, 4414–4423. URL: https://www.sciencedirect.com/science/article/pii/S0022030208709901.

Vitezica, Z. G., Legarra, A., Toro, M. A. and Varona, L. (2017) Orthogonal Estimates of Variances for Additive, Dominance, and Epistatic Effects in Populations. Genetics, 206, 1297–1307.

Vitezica, Z. G., Varona, L., Elsen, J.-M., Misztal, I., Herring, W. and Legarra, A. (2016) Genomic BLUP including additive and dominant variation in purebreds and F1 crossbreds, with an application in pigs. Genetics Selection Evolution, 48, 6. URL: 10.1186/s12711-016-0185-1.

Vitezica, Z. G., Varona, L. and Legarra, A. (2013) On the Additive and Dominant Variance and Covariance of Individuals Within the Genomic Selection Scope. Genetics, 195, 1223–1230. URL: 10.1534/genetics.113.155176.

Wickham, H. (2016) ggplot2: Elegant Graphics for Data Analysis. Springer-Verlag New York. URL: https://ggplot2.tidyverse.org.

Wilke, C. O. (2020) cowplot: Streamlined Plot Theme and Plot Annotations for ‘ggplot2’. URL: https://CRAN.R-project.org/package=cowplot. R package version 1.1.1.

Winn, Z. J., Amsberry, A. L., Haley, S. D., DeWitt, N. D. and Mason, R. E. (2023) Phenomic versus genomic prediction—a comparison of prediction accuracies for grain yield in hard winter wheat lines. The Plant Phenome Journal, 6, e20084.

Xiang, T., Christensen, O. F., Vitezica, Z. G. and Legarra, A. (2016) Genomic evaluation by including dominance effects and inbreeding depression for purebred and crossbred performance with an application in pigs. Genetics Selection Evolution, 48, 92. URL: 10.1186/s12711-016-0271-4.

Zhong, S., Dekkers, J. C. M., Fernando, R. L. and Jannink, J.-L. (2009) Factors Affecting Accuracy From Genomic Selection in Populations Derived From Multiple Inbred Lines: A Barley Case Study. Genetics, 182, 355–364. URL: 10.1534/genetics.108.098277.

